## Supplementary Information for "Role of non-specific interactions in the phase-separation and maturation of macromolecules"

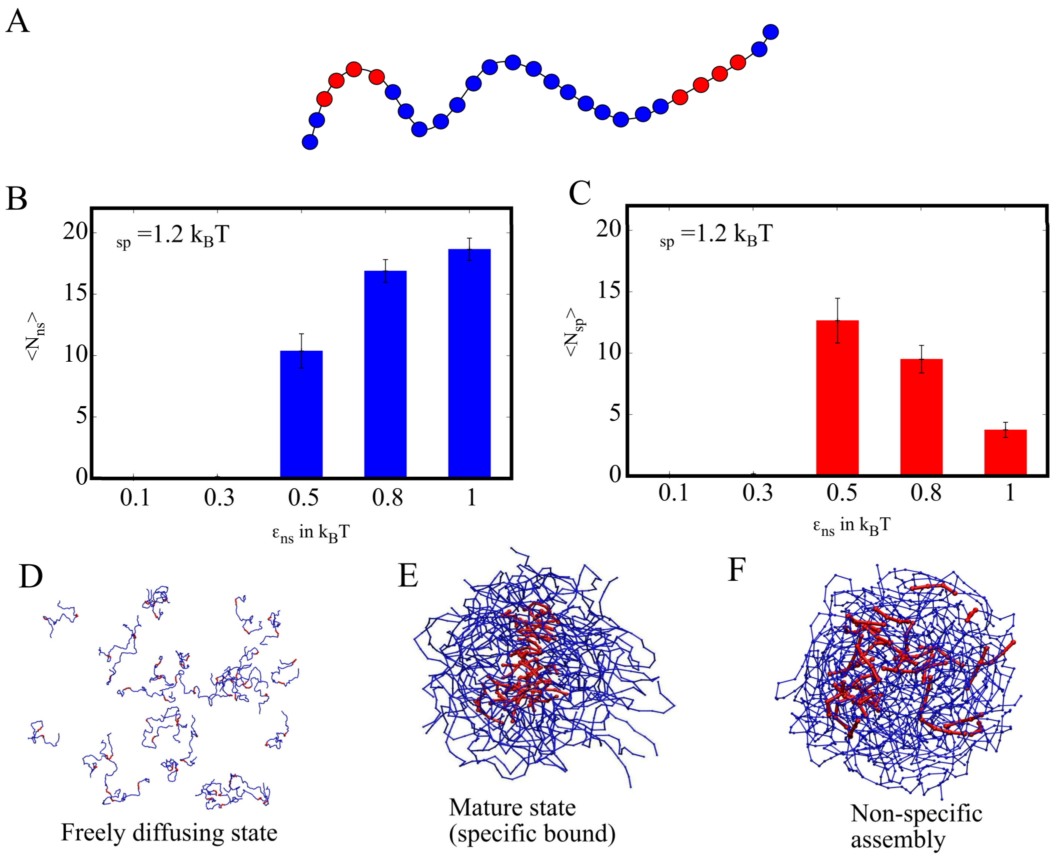


**Supplementary Fig. 1.** A) Bead spring polymer model of protein. Chain consisits of two patches of 4 red beads (specifically interacting) in the tail and other blue beads (non-specifically interacting) . B) Average number of non-specific contacts per bead (blue-blue and blue-red) for different $\epsilon_{\mathrm{ns}}$ values C) Average number of specific contacts per bead (red-red) for varying $\epsilon_{\mathrm{ns}}$. In for figure B and C, value of $\epsilon_{\mathrm{sp}} = 1.2$ $\mathrm{k_BT}$ is fixed. D) Snapshot of the simulation, where $\epsilon_{\mathrm{sp}} = 0.1$ and 0.3 $\mathrm{k_BT}$ – polymers are freely diffusing state. D) Snapshot of the simulation, where $\epsilon_{\mathrm{sp}} = 0.5$ and 0.8 $\mathrm{k_BT}$ – polymers are in mature state (specific/red-red bound). E) Snapshot of the simulation, where $\epsilon_{\mathrm{sp}} = 1.0$ $\mathrm{k_BT}$ – polymers are non-specifically bound.

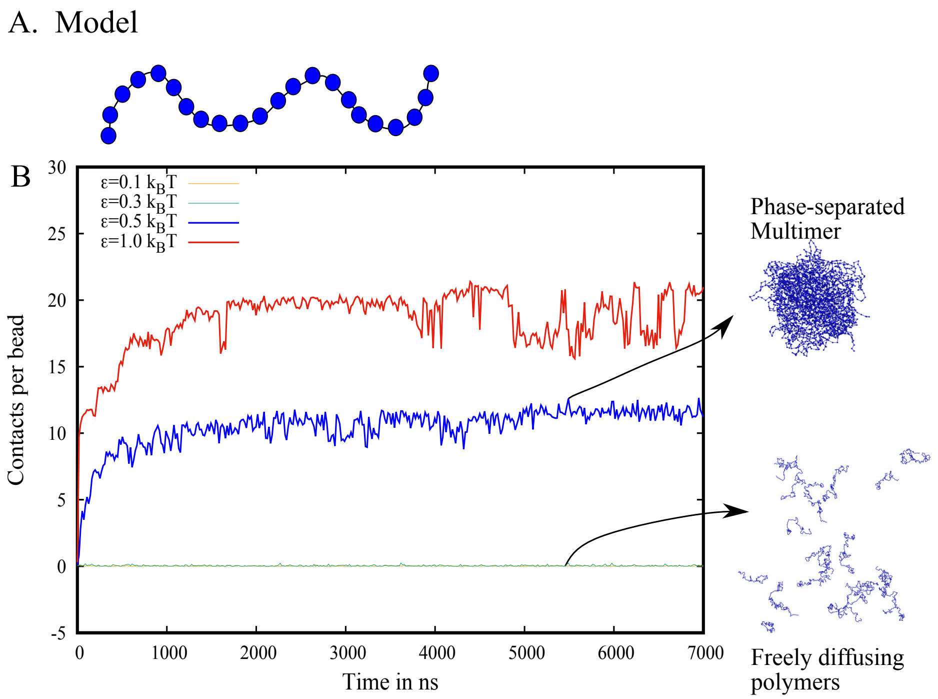


**Supplementary Fig. 2.** A) Bead spring polymer model of protein with all the beads in the polymer chain are made up same type of bead and have same interaction strength $\epsilon$. B) Contacts per bead over time is plotted for various values of interaction strength ($\epsilon$). Two different regimes of polymer phases, are observed namely freely diffusing polymers and phase-separated multimer.

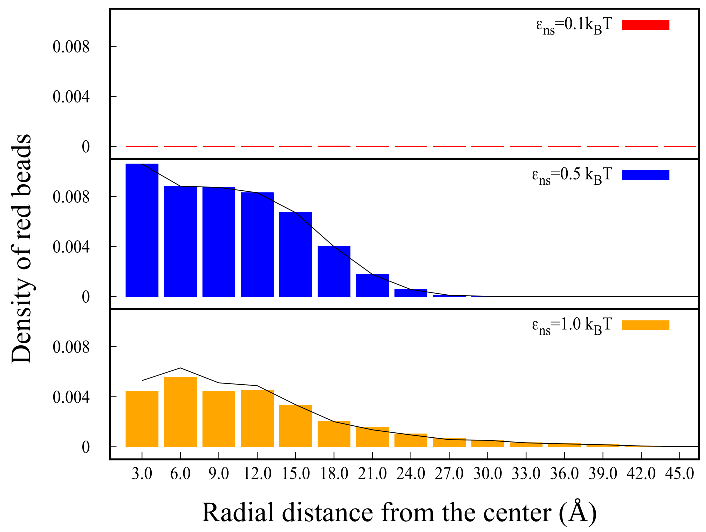


**Supplementary Fig. 3.** Density of red beads in each shell is plotted over the radial distance from the center.
